# Allele biased transcription factor binding across human brain regions gives mechanistic insight into eQTLs

**DOI:** 10.1101/2023.10.06.561245

**Authors:** Belle A. Moyers, Jacob M. Loupe, Stephanie A. Felker, James M.J. Lawlor, Ashlyn G. Anderson, Ivan Rodriguez-Nunez, William E. Bunney, Blynn G. Bunney, Preston M. Cartagena, Adolfo Sequeira, Stanley J. Watson, Huda Akil, Eric M. Mendenhall, Gregory M. Cooper, Richard M. Myers

## Abstract

Transcription Factors (TFs) influence gene expression by facilitating or disrupting the formation of transcription initiation machinery at particular genomic loci. Because genomic localization of TFs is in part driven by TF recognition of DNA sequence, variation in TF binding sites can disrupt TF-DNA associations and affect gene regulation. To identify variants that impact TF binding in human brain tissues, we quantified allele bias for 93 TFs analyzed with ChIP-seq experiments of multiple structural brain regions from two donors. Using graph genomes constructed from phased genomic sequence data, we compared ChIP-seq signal between alleles at heterozygous variants within each tissue sample from each donor. Comparison of results from different brain regions within donors and the same regions between donors provided measures of allele bias reproducibility. We identified thousands of DNA variants that show reproducible bias in ChIP-seq for at least one TF. We found that alleles that are rarer in the general population were more likely than common alleles to exhibit large biases, and more frequently led to reduced TF binding. Combining ChIP-seq with RNA-seq, we identified TF-allele interaction biases with RNA bias in a phased allele linked to 6,709 eQTL variants identified in GTEx data, 3,309 of which were found in neural contexts. Our results provide insights into the effects of both common and rare variation on gene regulation in the brain. These findings can facilitate mechanistic understanding of cis-regulatory variation associated with biological traits, including disease.

## Introduction

Gene expression changes occur in essentially every biological process, including the development of diseases (Emilsson et al. 2008; Lee and Young 2013) such as neurodegenerative (Bonham et al. 2019, 2022; Zhao 2023) and psychiatric conditions (Clifton et al. 2019; Mimmack et al. 2002; Huang et al. 2020). Transcription factors (TFs) and their association with DNA are crucial determinants of gene expression, so identifying factors that influence the association between TFs and DNA is key to understanding variation in gene expression. A wide variety of tools have been developed to identify and catalogue DNA sequence motifs to which TFs preferentially bind (Bailey et al. 2015; Ghandi et al. 2016; Castro-Mondragon et al. 2022). While informative, these approaches are limited by the fact that a motif’s presence is neither necessary nor sufficient for TF association (Dror et al. 2015), so the impact of DNA sequence changes on motifs is of limited utility.

An alternative approach is to leverage natural genetic diversity across and within humans, specifically heterozygous variants, and assay TF binding behavior. Tools have been developed to identify differential binding across multiple experiments to identify changes in TF binding (Lun and Smyth 2016), but these are complicated by technical and biological variation. Complicating this issue further is that reference allele bias is a known obstacle in mapping sequence reads, and this can inflate the false discovery rate in studies of allelic effects (Degner et al. 2009; Stevenson et al. 2013; Rozowsky et al. 2011; Smith et al. 2013; Hach et al. 2014). The use of graph structures to represent personalized genomes or pangenomes (Li et al. 2020; Paten et al. 2017) can reduce reference allele bias (Garrison et al. 2018; Martiniano et al. 2020; Chen et al. 2021).

A recent study probed allele-specific binding across hundreds of cell types with corrections for reference allele bias and aneuploid regions (Abramov et al. 2021). The majority of these datasets were derived from cancer cell lines, limiting their applicability to non-diseased human tissue, or in contexts relevant for specific disease states. Another recent study highlighted the viability of a similar approach in human tissue samples by identifying allele-specific loci with 15 assays in 4 human donors across 30 tissues, including ChIP-seq assays of histone marks and several TFs, including the CCCTC-binding factor CTCF (Rozowsky et al. 2023). This study found relationships between allele-specificity of ChIP-seq and gene expression, including identifying GTEx eQTLs that were allele-specific and those that were not. This highlights the value of identifying allele-biased binding among transcription factors for understanding gene regulation.

Here, we greatly expand upon previous work by performing allele-biased binding analysis for 1,004 (Loupe et al. 2023) TF-ChIP-seq datasets, spanning 93 distinct TFs, RNA polymerase II (POLR2), and 5 histone marks in tissue samples from 9 anatomically defined brain regions in multiple donors. We used the vg toolkit to assemble personalized genomes to overcome reference allele bias and demonstrated that this approach improves the calling of allele-biased binding. We explored dynamics of allele frequency in the population with allele bias and the relationship between allele-biased binding and disruption of TF binding motifs. We determined the effects on gene expression by using RNA-seq reads to assess eQTLs in these donors, allowing a mechanistic exploration of eQTL data. Finally, we highlight interesting examples of allele-biased binding identified in our datasets.

## Results

### Graph Genomes improve read mapping and reduce reference allele bias

To study the impact of genetic variation on TF binding, we first performed linked-read sequencing (10x Genomics) to generate phased genomes and call variants for two donors (**Figure 1A**; see Methods). We built personalized graph genomes that use the vg toolkit (Garrison et al. 2018), as it has been shown that it can reduce problems of reference allele bias and has previously been used for detection of missing signal in histones (Groza et al. 2020) and the detection of allele-biased TF footprints (Ouyang and Boyle 2022). To measure the effectiveness of this approach on our datasets, we initially mapped a pilot set of 20 ChIP-seq datasets, 10 in each of the two donors, using both conventional mapping to the hg38 linear reference via bowtie2 and personalized graph-genome mapping via vg. We observed an average increase in read mapping of 1.24% of the total read pool when using personalized graph genomes (**Table 1**). Despite this small change in overall mapping, the use of graph genomes greatly reduced the degree of reference allele bias for variants identified as significantly biased using the two approaches (**Figure 1B**). Because rare alternate alleles tend to be deleterious and may show increased preference for the reference allele, we restricted to cases of MAF>0.05 for this analysis. Using hg38, we identified 21,207 cases of significant TF-allele bias at the nominal p<=0.05 level (binomial test), 76.9% of which favor the reference allele, compared to 17,823 cases of significant TF-allele bias using a graph genome, with 52.8% favoring the reference allele. The reference bias trend was also observed, though reduced, when considering all variants, including those with MAF<=0.05 with bias (**Supplemental Figure 1**), as well as variants with at least six mapped reads whether or not there was a nominally significant bias (**Supplemental Figure 2**). Thus, graph genome alignments tend to reduce reference-alignment artifact contributions to observed allelic biases.

**Figure 1.**
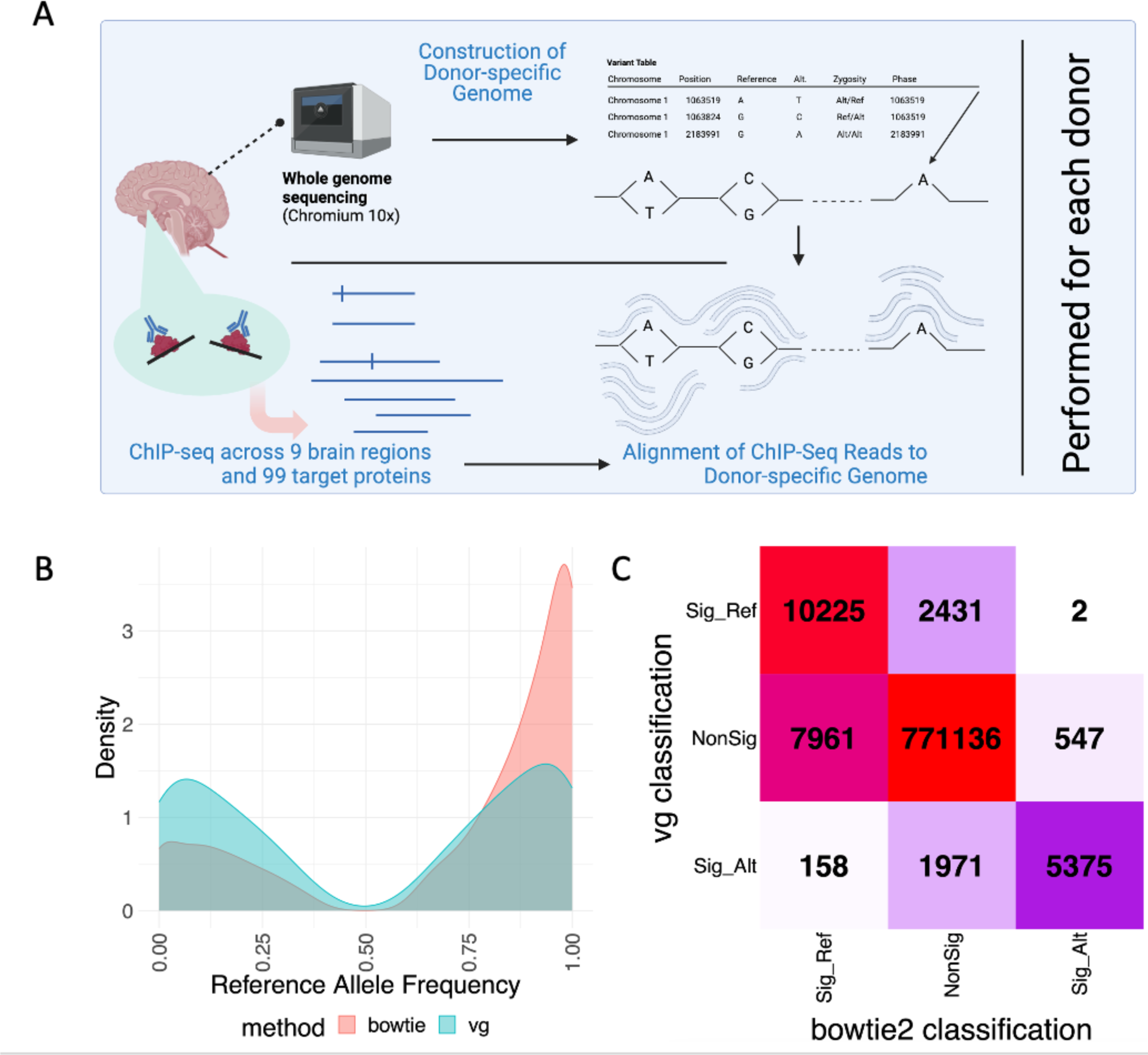
Personalized graph genomes improve read mapping for detection of allele-biased binding. **A**. Workflow for detection of allele biased binding. Whole Genome Sequencing and ChIP-seq of 93 TFs, POLR2, and 5 histone marks were performed in post-mortem brain samples from 2 donors. ChIP-seq reads were mapped to personalized graph genomes to identify allele bias and were compared within and across donors. **B.** Personalized genomes reduce problems of reference allele bias, increasing confidence in allele-biased binding detection. Density plots are shown for the reference allele frequency (x-axis) of significant (*p*<=0.05, binomial test) allele bias when using bowtie aligned to the linear reference (red) compared to using the vg toolkit aligned to a personalized graph genome (blue). Allele bias is more balanced between the reference and alternate for personalized graph genomes. **C**. There is significant disagreement in the number and identity of variants found preferring the reference and alternate alleles between methods. Heatmap showing the number of TF-biased allele interactions found nonsignificant, significant for reference, and significant for alternate by bowtie and vg.

**Table 1.** Percentage of reads mapped generally increases using personalized genomes compared to linear genomes. For 20 ChIP-seq datasets, the percentage of reads which were mapped to the reference genome when using a linear genome (Linear) or a personalized graph genome (Graph) for donor 1 (left) and donor 2 (right). In most cases, vg maps a larger percentage of reads.

|  | Percent of reads Mapped |  |  |  |
| --- | --- | --- | --- | --- |
|  | Donor 1 |  | Donor 2 |  |
|  | Linear | Graph | Linear | Graph |
| **Control** | 99.48 | 99.95 | 99.47 | 99.95 |
| **BCL11A, DLPFC** | 86.88 | 91.61 | 90.69 | 93.23 |
| **CREB1, CB** | 98.54 | 99.37 | 98.65 | 98.24 |
| **CTCF, OL** | 94.99 | 96.97 | 94.72 | 96.15 |
| **CTCF, SG** | 96.24 | 97.70 | 96.30 | 96.86 |
| **MEF2A, DLPFC** | 97.21 | 98.47 | 96.83 | 97.22 |
| **RAD21, ANCG** | 96.64 | 98.04 | 97.62 | 97.60 |
| **RAD21, HC** | 95.80 | 97.46 | 96.77 | 96.97 |
| **SP1, AMY** | 94.35 | 96.49 | 93.37 | 95.15 |
| **TBR1, DLPFC** | 96.41 | 97.93 | 96.65 | 96.91 |

To determine how specific variants were classified by each method, we next created a heatmap of the number of variants classified as significant or nonsignificant with each mapping method (**Figure 1C**), and which of the alleles—reference or alternate—they preferred. We found that, for TF-allele interactions identified as significant in both methods, the direction of effect is well-conserved between the two methods. However, we found 4,402 variants that were significant only when we used a graph-based approach, and 8,508 variants that were significant only when we used a linear reference. For these variants, we note that bowtie2’s mapping produces a strong preference for predicting the reference allele as the preferred allele, with 7,961 of the 8,508 hg38-specific allele biased events favoring the reference. In contrast, vg shows a more balanced distribution of variants favoring the reference and alternate alleles (55.2% favoring reference). This is consistent with the problems of reference allele bias seen in **Figure 1B**. In addition, when the two methods disagree in their classification of a variant, vg tends to have a higher read depth at the location (**Supplemental Figure 3**), suggesting that improved mapping of reads with variants results in a change in the apparent significance of the variant. Together, these findings suggest that use of personalized genomes substantially improves both specificity and sensitivity for detection of TF-allele bias.

### Allele-biased binding is consistent across donors and tissues

We subsequently measured TF allelic bias using only the graph genome approach, applying it to ChIP-seq data from 93 TFs, RNA Polymerase (POLR2A), and five histone marks in up to nine anatomically defined regions of the brain across two donors, for a total of 1,004 ChIP-seq datasets (Loupe et al. 2023). We used vg to map these ChIP-seq datasets and calculate allele bias for each dataset in each haplotype. We initially identified all allele bias at a nominal p-value <= 0.05 (binomial test). At that threshold, we found that of the nearly two million regions with heterozygous DNA sequence variation in each donor, roughly 7.5% were significantly biased for at least one ChIP-seq dataset (**Table 2**); as expected, this largely reflects the fact that TF binding occurs at only a small fraction (7-10%) of genomic loci (Loupe et al. 2023). We note that, while 266,448 variants are heterozygous in both donors (13.8%-14.1% of all heterozygous sites in each donor), only 5,954 heterozygous variants that showed significant bias in either donor are significantly biased in both donors (4.1-4.2%). This points to a 3-fold depletion of shared TF-biased variation between our two donors, suggesting that selection reduces the frequency of such variation in the population (*p*<=2.2 x 10^-16^, binomial test).

**Table 2.** The number of variants found significant in each donor individual, as well as the shared set of variants. When parentheses are present, the number outside of the parentheses denotes the number of variants found significant in at least one of the two donors, while the number inside the parentheses shows the number which were significant in both donors. Percentages denote the percent of this intersect (within parentheses) compared to the union (outside of the parentheses).

|  | **Donor 1** | **Donor 2** | **Shared**<br>**(Significant in**<br>**Both Donors)** |
| --- | --- | --- | --- |
| Variant regions | 3,032,858 | 3,074,271 | 809,297 |
| Heterozygous regions | 1,894,277 | 1,932,258 | 266,488 |
| Significantly Biased<br>($p \leq 0.05$) | 139,234 | 144,952 | 28,751 (5,862;<br>20.4%) |
| Significantly Biased<br>($p \leq 0.001$) | 4,328 | 3,876 | 486 (142; 29.2%) |
| Significantly Biased<br>when summed<br>across tissues<br>($p \leq 0.001$) | 7,828 | 9,570 | 1,195 (377; 31.5%) |

We next assessed reproducibility. Given that we performed experiments in tissues from multiple brain regions within each donor and two donors for each, we assessed consistency of results both on the same region between the two donors and between different regions within the same donor. Each comparison type captures a different mixture of technical and biological factors. Cross-donor/within-region differences may be due to either experimental errors or genuine between-donor differences, while cross-region/within-donor differences may be due to experimental errors or genuine region-level differences.

We first assessed between-donor reproducibility of the effects of shared variants by determining whether or not the variant was consistent in its effect direction between the two donors. For each shared variant (i.e., both donors are heterozygous) that was significant in at least one donor for a given TF in a given region, we determined the number of reads mapping to each allele in both donors (restricting to cases with at least six total reads mapped) and determined whether or not the same allele is favored in both donors. We measured effect direction reproducibility as 100% minus twice the percentage of inconsistent effect direction observations, as half of all random comparisons would by chance appear to be consistent (i.e., if 10% of comparisons exhibit inconsistent effect directions, the inferred reproducibility rate is 80%). We then assessed reproducibility across a range of nominal bias p-value thresholds (**Figure 2A**). We found that at a p-value cutoff of 0.05, just under 60% of variant effects on TF binding are inferred to be reproducible across donors, but that at a cutoff of 0.001 that increases to more than 85% of the variants.

**Figure 2.**
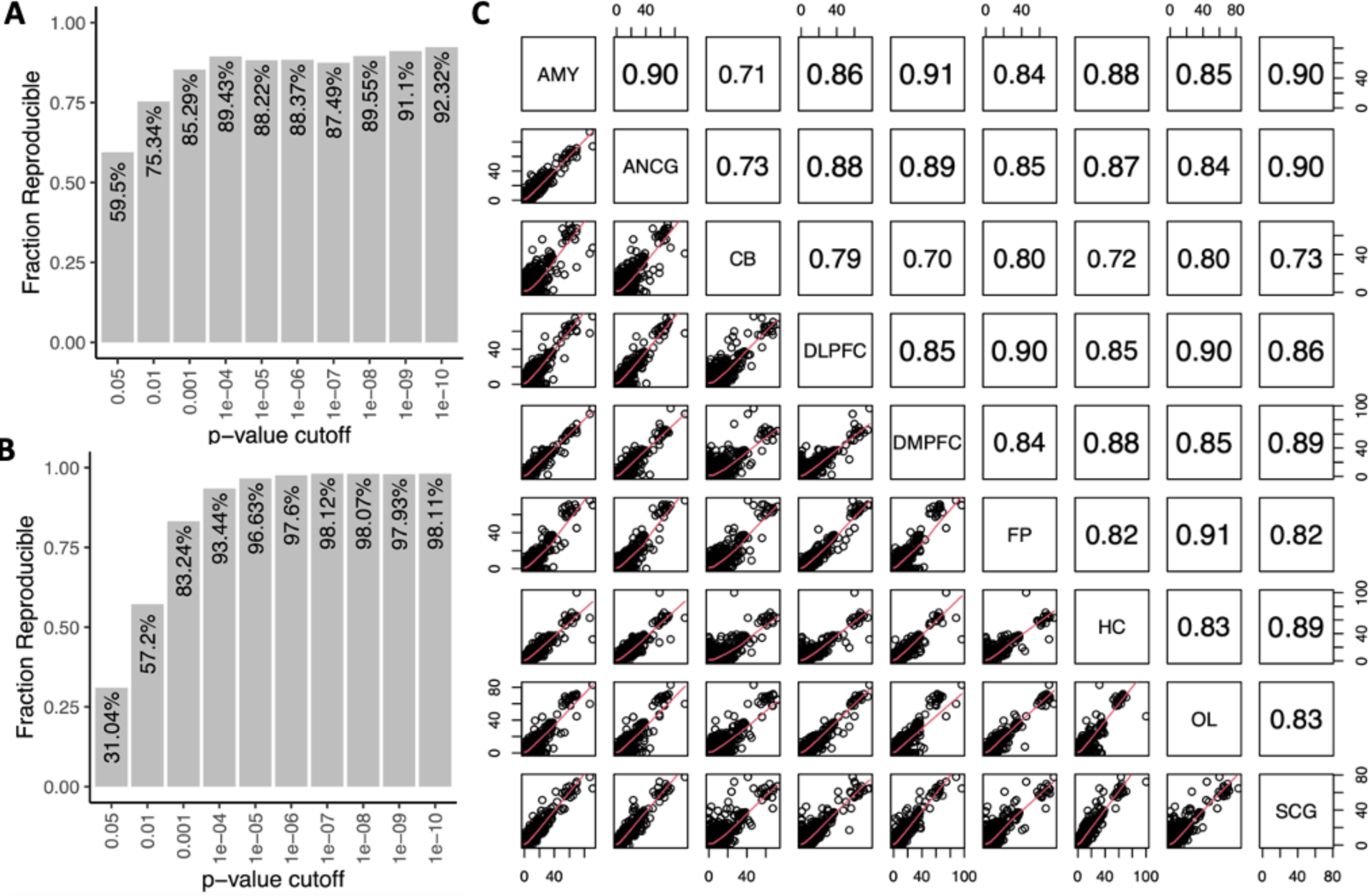
Reproducibility and concordance of TF-allele bias within and between donors. **A.** Between-donor reproducibility. The fraction of TF-allele bias cases which were reproducible in the comparable TF-allele interaction in the same tissue across donors (y-axis) as a function of the minimum p-value cutoff used for significance (x-axis). Reproducibility was defined as 1 – 2*(Percent of inconsistent directional effects identified). **B.** Within-donor reproducibility. The fraction of TF-allele bias cases which were reproducible when comparing the same TF-allele interaction across different tissue contexts within the same donor. Reproducibility was defined as in 2A. **C**. Correlation of −log(p-value) of effects of a variant across tissues, for variants with a pvalue of <=0.001 in at least one tissue for factors with ChIP-seq datasets in all 9 tissues. Bottom shows dot-plots of variant effects. Top shows correlation coefficients (Pearson) between each tissue. Diagonal line notes each tissue. Abbreviations denote: dorsolateral prefrontal cortex (DLPFC), frontal pole (FP), occipital lobe (OL), cerebellum (CB), anterior cingulate (AnCg), subgenual cingulate (SCg), dorsomedial prefrontal cortex (DMPFC), amygdala (Amy), and hippocampus (HC).

We also explored within-donor reproducibility between regions. This analysis yielded far more comparisons, as all heterozygous variants are shared across all regions within the same donor, and each TF could be compared across up to nine brain regions, resulting in up to 36 total comparisons for each variant’s impact on TF binding. We therefore assessed within-donor reproducibility in the following way. We determined all variants that impacted a TF’s binding in at least one brain region. We then looked in each brain region where data were available for that TF with at least six total mapped reads and counted the number of reads mapping to each allele. We then determined reproducibility as described above, at each p-value cutoff. We found that at a p-value cutoff of 0.05, reproducibility is only marginal at >30%, but that a p-value cutoff of 0.001 between-region reproducibility was more than 80% (**Figure 2B**).

Based on these observations, we restricted further analyses of significant variants to those with a nominal p-value <= 0.001 for a reproducibility rate of >80% unless otherwise noted. This metric confidently identifies allele-biased TF-DNA interactions both within and across donors. A summary of the number of variants impacting TF binding at this cutoff is included in **Table 2**.

Because within-donor reproducibility was high, we assessed the overall correlation across brain regions simultaneously for TF-DNA interactions. The ChIP-seq datasets we analyzed fall into two categories, four large brain regions (cerebellum, dorsolateral prefrontal cortex, occipital lobe and frontal pole), which provided enough material to do ChIP-seq on 93 TFs, and five smaller brain regions, which provided enough material for only 16 TF ChIP-seq maps. For each pairwise comparison of brain regions, we determined the correlation of the −log10(pvalue) for each TF-DNA biased interaction we identified (**Figure 2C**). We note that correlations between tissues range between 0.70 and 0.91 (Pearson’s correlation coefficient) when using TFs with ChIP-seq data (n=16) in all nine brain regions, and 0.81-0.91 when using a larger number of TFs (n=93) limited to four brain regions (**Supplemental Figure 4**). The cerebellum showed good but comparatively lower correlation with other tissues. This is expected, as the cerebellum has markedly different cellular makeup than other brain regions (Andersen et al. 1992; Loupe et al. 2023). Overall, this analysis shows a strong quantitative correlation for TF-variant bias across multiple regions of the brain.

Because variants have strong reproducibility across and within donors and have high correlation in their effect size of impact upon TF association within donors, we combined reads across all brain regions for each variant for a given ChIP-seq target for other downstream analyses (**Table 2**). Given the extra statistical power from combining reads, we observed a 2-fold increase in the identified TF-biased variants at p<=0.001: 7,828 TF-biased variants (0.41%) in Donor 1 and 9,570 (0.50%) in Donor 2 (at *p*<=0.001). Among these variants, we asked how many showed corroborating bias in POLR2A or any of the histone datasets, which would not be predicted to be directly altered by variants, but are likely to reflect altered gene regulation at that variant. We identified those variants that were also biased for at least one histone mark or POLR2A, and found that approximately half are biased for at least one of these datasets (4,271 in donor 1 and 4,734 in donor 2). We found that, for such variants, more TFs are generally biased for the variant (**Supplemental Figure 5**), and that the significance of TF bias increases (**Supplemental Figure 6**).

**Supplemental Tables 1 and 2** show all nominally significant (p<=0.05) heterozygous regions for any ChIP-seq target or input DNA in each donor based upon summed reads across brain regions as well as relevant information about each variant. Hereafter, we consider only those variants that impact TF binding, independent of POL2RA or histone effects, at p<=0.001.

### Allele Bias is prevalent in functional regions important for neuronal differentiation

We next explored the genomic properties of the 17,309 unique variants that impact TF binding using candidate Cis-Regulatory Elements (cCREs) from the ENCODE Consortium (Luo et al. 2020), which marks elements such as promoters and enhancers. We classified each heterozygous region, TF ChIP-seq peak, and allele-biased variant into cCRE categories (**Figure 3A**). Not surprisingly, we found that ChIP-seq peaks overwhelmingly lie within cCRE regions, while the majority of heterozygous variation falls outside of cCRE regions. However, most cases (73.3%) of TF-allele bias fall within or near cCREs, despite requiring only a minimum of 11 reads total across all experiments to potentially be found as significantly biased at p<=0.001 for a given variant. Despite being overwhelmingly within cCREs, 83.2% of allele-biased variants are not in a called peak for that TF.

**Figure 3.**
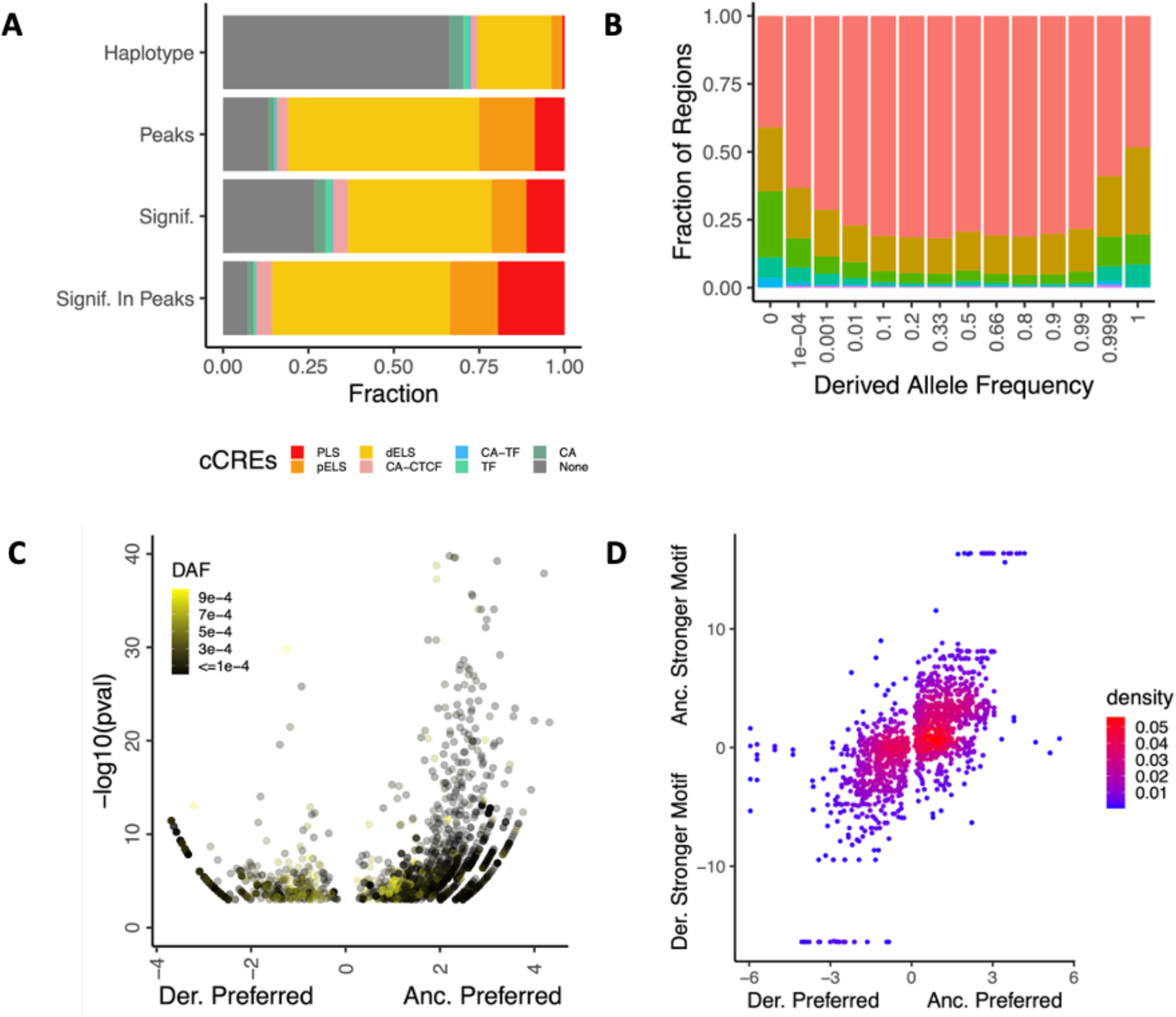
Genetic and genomic properties of variants displaying TF-allele bias. **A.** Stacked barplots showing the fraction of regions which have overlap with a particular cCRE type for all variant haplotypes (first from top), all TF peaks (second from top), haplotypes found significant for at least one TF (second from bottom), and haplotypes found significant in at least one TF while also overlapping with a TF peak (bottom) (y-axis). Cumulative fraction is shown on the x-axis. Barplots are colored by cCRE type as PLS (promoter-like signal): red, pELS (proximal enhancer-like signal) orange, dELS (distal enhancer-like signal) yellow, CA-CTCF (chromatin-accessible CTCF signal) pink, CA-TF (chromatin-accessible, TF signal) blue, TF (TF signal) blue-green, CA (chromatin-accessible) green, and with non-cCRE regions plotted in grey. **B.** Variants which are either very rare or very common in the population show highly significant allele bias. For varying ranges of derived allele frequency (x-axis), we show the fraction of significant variants which were found at or below a given significance threashold (y-axis). **C.** For very low-frequency derived alleles, a volcano-like plot is shown which relates the ChIP-seq preference for the ancestral allele (x-axis, log(ancestral ChIP-seq reads+1 / derived ChIP-seq reads+1) and the significance (y-axis, −log10(pvalue) as determined by a binomial test) for each significantly-biased variant. Points are colored by their derived allele frequency, with rarer derived alleles being black and more common, up to DAF=0.001, being plotted in yellow. For very rare alleles, there is a stronger preference for the ancestral allele, and the significance of bias is higher. **D.** For variants which weaken or strengthen a JASPAR motif (i.e. a motif was found in each sequence, but the score changed) for one of our assays TFs, the difference in FIMO score between the ancestral and derived allele (y-axis) versus the log(ancestralReads/derivedReads) for the relevant TF. Spearman’s Rho = 0.658, p<=2. X 10^-16^.

Restricting to only those cases of allele-biased binding within peaks, we found that, relative to global peak locations, allele-biased binding is 1.8-fold enriched for PLS regions (*p*<=2.2 x 10^-16^, binomial test). We also found that a substantial number of biased variants occur in dELS and pELS regions, consistent with noted trends in the evolution of variability in enhancer function over evolutionary time (Rebeiz and Tsiantis 2017; Lynch et al. 2015; Emera et al. 2016).

To explore the function of these regions, we performed GO analysis using GREAT (McLean et al. 2010; Gu and Hübschmann 2023), using allele-biased regions as our regions of interest and all peak regions minus allele-biased regions as controls (**Supplemental Table 3**). We note that the top 10 enriched terms (sorted by adjusted hypergeometric p-value) are largely involved in neural development, organismal development, and cell communication.

### Very rare variants are more likely to disrupt TF-DNA associations

Because allele-biased regions are found near neural and developmental genes, suggesting a functional outcome on cellular and developmental phenotypes, we explored how common these variants are in population databases. We hypothesized that any derived allele which significantly altered the expression of a key neural or developmental gene would experience natural selection during human evolution. We identified ancestral alleles via comparative genomics among apes using Ensembl (Cunningham et al. 2022) and used gnomAD (Chen et al. 2022) to identify current allele frequency in the human population. We then binned TF-biased variants by derived allele frequency and noted the fraction in each bin with a given allele bias p-value (**Figure 3B**). We found that, for rare variation in the population (extreme right and left derived allele frequency bins), bias tends to be more significant than for common variation (middle bins). However, we note that there are relatively few allele-biased variants that are rare in the population but for which the derived allele is more common (e.g., 41 variants with DAF >0.999, vs 719 with DAF < 0.001).

To further analyze effects of the ancestral and derived alleles among rare variants, we selected the TF-biased variants with derived allele frequency (DAF) of 0.001 or lower (791 variants) and determined the number of reads that map to the ancestral and derived alleles (**Figure 3C**). We found a strong preference for the ancestral allele both in the number of cases supporting it (70.2% support ancestral versus 29.8% derived), and degree of bias significance (**Figure 3C**). When restricting to variants with very high DAF (>=0.999) (41 variants), we observed the opposite bias (33.7% support ancestral, 66.3% support derived), (**Supplementary Figure 7**). Among variation with DAF between 0.001 and 0.999 (i.e., sites at which both alleles are frequently observed in the human population, 24,818 variants), there is much reduced ancestral versus derived bias (56.2% vs 43.8%, **Supplemental Figure 8**). This suggests that common alleles are approximately equally likely to increase or decrease TF-DNA associations, whereas rare alleles are more likely to specifically disrupt TF-DNA association, while a smaller fraction appear to lead to new TF-DNA associations.

To evaluate the mechanism of allele-biased variation on TF-binding, we identified motifs that are disrupted by a heterozygous variant using human motifs for relevant TFs in the JASPAR database (Castro-Mondragon et al. 2022), and the fimo function of the meme (Bailey et al. 2015) suite. We first checked, for each TF, each biased variant and asked what percentage of the time the motif for that TF was significantly disrupted from consensus. We found a wide range for this metric, with 0-44% of the biased loci showing disrupted motifs. This likely reflects each TF’s motif strength. For example, the zinc finger factor CTCF, which has a long 14 bp consensus motif with many highly conserved bases, had its motif disrupted at 44% of the loci showing bias for that factor. By comparison, MAZ, which has a 7 bp motif with no strongly conserved nucleotides, had a disrupted motif in only 9.5% of TF-biased loci (**Supplemental Figure 9, Supplemental Table 4**). We next identified cases where a motif’s score changed between the two alleles and determined whether the derived or ancestral allele had a higher score, and the number of reads mapping to each allele. We found that allelic disruption of a motif is moderately correlated (Spearman’s Rho = 0.658) with TF ChIP-seq reads mapped (**Figure 3D**). This is also true of variants that entirely remove or create a motif, defined as finding a fimo hit in one allele and none at all in the other (**Supplemental Figure 10**) (Spearman’s Rho = 0.494).

Because we observed these trends in enrichment and in motif modifications, we then determined whether or not there was evidence for enrichment or depletion of sites with TF-biased variation being under purifying selection throughout mammalian evolution. We used the Genomic Evolutionary Rate Profiling (GERP) (Davydov et al. 2010) metric and identified those variants with GERP>4, a commonly-used cutoff for selective constraint (Schubert et al. 2014; Marsden et al. 2016). For each TF, we identified all variant locations with at least 11 reads mapped (minimum number of reads for binomial significance of 0.001), and determined the number of variants with GERP>4 and with GERP<4 for variants that were significant and for those that were non-significant. (**Supplemental Figure 11**). We find that, while most TFs have an apparent depletion of biased variants under selective constraint, none are significantly depleted (Chi-squared test).

### Allele-biased binding offers insight into eQTL mechanisms

Because TFs regulate the expression of RNA, we explored the relationship between allele-biased binding and the GTEx (GTEx Consortium 2020) database of expression quantitative trait loci (eQTLs) (Nica and Dermitzakis 2013). We found that 51.98% of all significant GTEx variants were present in at least one of our donors in either a heterozygous or homozygous state. We found 1,142,111 variants in a heterozygous state in Donor 1, and 1,123,497 in Donor 2, a necessary condition for detecting allele-biased binding of TFs. Of these variants, we found significant TF-allele bias in 7,459 variants in Donor 1 and 7,975 in Donor 2 for a total of 14,419 unique variants. We found that the involvement of allele-bias for individual TFs in GTEx eQTLs is similar to the genome at large (**Supplemental Figures 9 and 12**).

We next explored RNA-seq allele bias by mapping total RNA-seq reads to personalized genomes, determining the number of reads preferring each allele, summing across tissues, and identifying variants with a bias p-value <= 0.001. Using a model of known genes in the hg38 build (Bioconductor Core Team 2017), we determined which of these cases of allele bias overlapped with known genes. In Donor 1, 80.86% (5,850 of 7,191 total) of allele-biased RNA reads occurred in known gene models, and 81.35% (5,774 of 7,141 total) in Donor 2. We detected allele-biased expression in 10.16% of gene bodies in Donor 1 and 11.32% in Donor 2, consistent with estimates of the fraction of genes with allele-biased expression in earlier studies (Gimelbrant et al. 2007; Kravitz et al. 2023).

We next identified variants in eQTLs that displayed both TF-allele bias as well as in-phase allele-biased RNA expression within the appropriate gene body as noted in GTEx. We found 6,709 GTEx variants that existed in a heterozygous state in one or both of our donors and with both a ChIP-seq allele bias and an in-phase heterozygous variant in the appropriate gene body with an RNA-seq allele bias. Because eQTLs can be tissue-specific (Mizuno and Okada 2019), we restricted to GTEx variants with annotations in neural tissue for further investigation. We found 3,309 of these variants were identified in the brain or neural tissue by GTEx. For each of these 3,309 variants, we identified the predicted slope of the variant for a given gene as well as the degree and direction of observed RNA-seq allele bias in our reads. We found a modest, but highly significant, correlation of 0.43 (Spearman’s Rho, *p*<=2.2 x 10^-16^) (**Figure 4A**). This suggests a mechanistic link between allele-biased TF binding and RNA expression, consistent with the general function of TFs, that at least partially explains population-wide genetically-determined expression variation.

**Figure 4.**
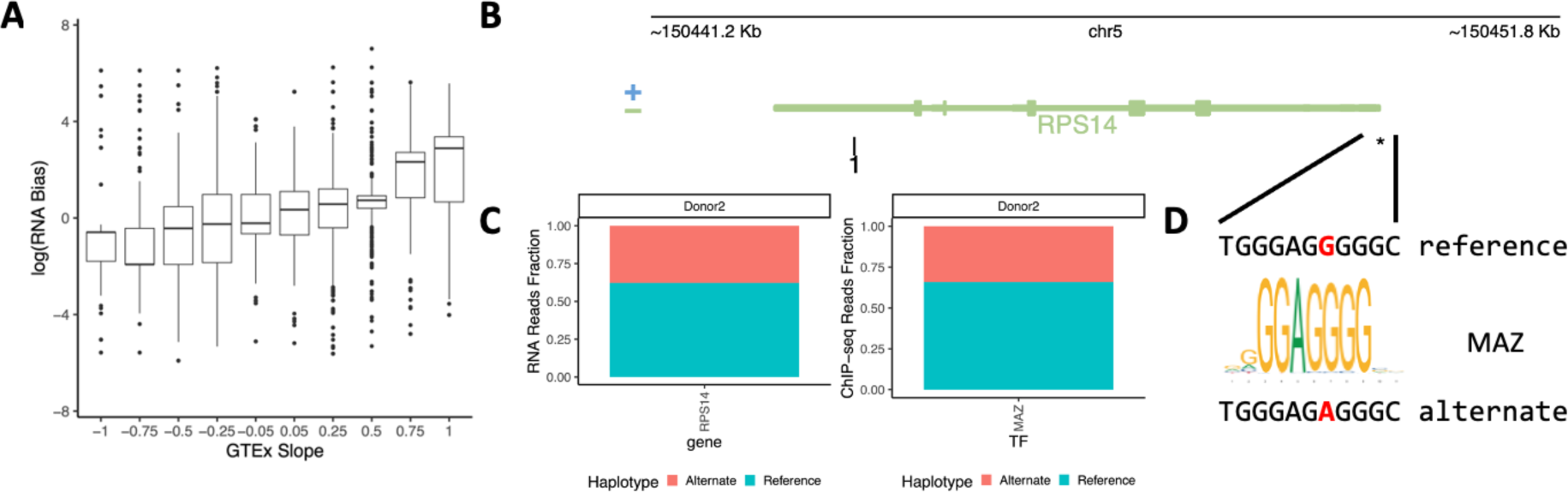
Allele-biased binding is consistent with and offers insight to the mechanisms of GTEx eQTLs. **A**. For GTEx eQTLs found in a neural context present in our data with significantly-biased ChIP-seq signal and phased significantly-biased RNA-seq reads in the appropriate genic region, a violin plot showing the distribution of log(RNA bias) (y-axis) versus the binned GTEx eQTL slope (x-axis). Spearman’s Rho 0.43, p<=2.2 x 10^-16^. **B.** Genomic track for the RPS14 gene showing the location of the GTEx eQTL chr5_150449748_G_A_b38 in the promoter. Green genes represent presence on the reverse strand, blue genes represent presence on the forward strand. Asterisk denotes the position of the eQTL. Tick marks denote heterozygous variants in the same phase as our heterozygous eQTL. **C.** Stacked barplots showing the fraction of reads supporting the reference or alternate strand (y-axis) of the eQTL for RNA (left) or ChIP-seq reads for biased TFs (right). **D.** Sequence of DNA surrounding the eQTL in B for the reference (top) and alternate (bottom) alleles, with the eQTL variant highlighted in red. Between them is displayed the MAZ motif MA1522.1 found in JASPAR, highlighting the alternate allele’s destruction of the canonical motif.

We highlight a simple case found in Donor 2, chr5_150449748_G_A_b38, in **Figure 4B-D**. The variant occurs near the TSS of the *RPS14* gene (**Figure 4B**), which encodes a ribosomal protein. This eQTL was found to be significant in 10 tissue contexts in GTEx, with slope values from −0.27 to −0.11, meaning that the alternate allele decreases expression relative to the reference allele. We found that RNA expression in our dataset is biased in the expected direction (**Figure 4C, left**), and that there is TF-allele bias over the variant for the MAZ transcription factor (**Figure 4C, right**). Comparing the two sequences, we find that the alternate variant disrupts the MA1522.1 motif of the MAZ (**Figure 4D**).

In another case, we explored a more complicated eQTL case found in a heterozygous state in both donors, chr22_32474782_C_T_b38, in **Figure 5**. This variant occurs in the promoter region of *FBXO7* (**Figure 5A**), an F-Box protein with a suggested role in Parkinson’s Disease (Joseph et al. 2018; Conedera et al. 2016; Burchell et al. 2013). We found that Hi-C data from iCell GlutaNeurons (Rogers et al. 2023) supports this locus interacting with several distal regions (**Figure 5A**). This variant was found in an eQTL for *FBXO7* expression in eight tissues with slope values from 0.2 to .45. We found that our RNA-allele bias data also supports the alternate allele having a higher expression (**Figure 5B, left**), and that several TFs in each donor also prefer the alternate allele (**Figure 5B, right**). Interestingly, this variant is also found to be associated in GTEx with a tissue-dependent change of expression of the *SYN3* gene, a neuronal phosphoprotein that associates with the surface of synaptic vesicles. We had limited ability to detect such changes, with only a single heterozygous RNA-biased variant in Donor 2 in phase with the eQTL variant, but found a strong preference for expression of the *SYN3* reference allele. The fact that this variant occurs in a known CTCF binding motif (MA0139.1) (**Figure 5C**) and shows TF-allele bias for cohesion factors (**Figure 5B, right**) suggests some measure of distal action for this variant consistent with CTCF’s known roles (Splinter et al. 2006). We also observed several other phased variants were present in our donors in this region, each for the *FBXO7* gene. We explored allele-biased binding at these variants and highlight our findings in **Table 3**. Of note, several of these variants show no bias for any of our tested transcription factors or histone marks. This suggests that allele-biased binding may be a method of fine-mapping eQTLs when data are available.

**Figure 5.**
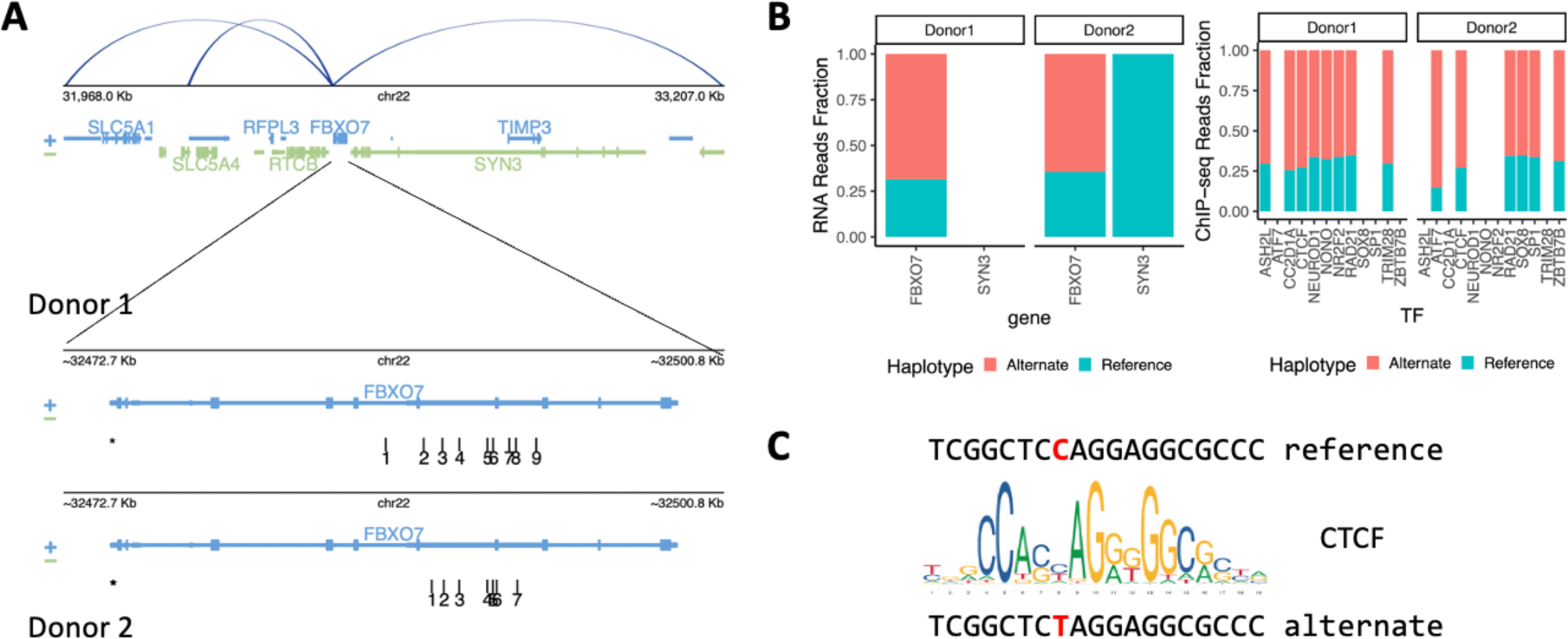
Allele-biased binding allows for fine-mapping of eQTL variants. **A**. Genomic track showing the region surrounding the GTEx eQTL chr22_32474782_C_T_b38, found in heterozygous form in both donors. Green genes represent presence on the reverse strand, blue genes represent presence on the forward strand. Asterisk denotes the position of the eQTL. Tick marks denote heterozygous variants in the same phase as our heterozygous eQTL. Loops from Hi-C data in iCell GlutaNeurons are shown above the gene tracks, noting 3D interactions. **B**. Left: Barplots depicting the fraction of reads supporting the strand of the reference (blue) or alternate (red) strand with regard to the eQTL for donor 1 and donor 2 for each of the FBXO7 or SYN3 gene. Right: Barplots depicting the fraction of reads mapping to the reference or alternate allele of the eQTL for significantly biased cohesion factors in donor 1 and donor 2. **C.** Sequence of DNA surrounding the eQTL in **A** for the reference (top) and alternate (bottom) alleles, with the eQTL variant highlighted in red. Between them is displayed the CTCF motif MA0139.1 found in JASPAR, highlighting the variant site.

**Table 3.** For variants found in GTEx which were in-phase with the focal variant, chr22_32474782_C_T_b38 (marked with *, explored in Figure 5), we show whether or not the variant was present in each of the two donors, as well as note any TF with significant allele-biased binding for that variant. We note that chr22_32474819_C_T_b38 (marked with **) was found within 100bp of the focal eQTL in donor 2, and so was analyzed in conjunction with the focal eQTL (see methods).

| GTEx Variant | Donor 1 Presence | Donor 2 Presence | Donor 1 Significant ChIP-seq | Donor 2 Significant ChIP-seq |
| --- | --- | --- | --- | --- |
| chr22_32470947_T_C_b38 | Yes | Yes | None | None |
| chr22_32471256_A_T_b38 | No | Yes | NA | None |
| chr22_32471173_G_C_b38 | Yes | Yes | None | None |
| chr22_32471541_G_A_b38 | Yes | Yes | None | None |
| chr22_32473024_C_T_b38 | Yes | Yes | None | None |
| chr22_32473508_G_T_b38 | Yes | Yes | CTCF | CTCF |
| chr22_32474674_C_T_b38 | Yes | Yes | None | POL2, ATF7,<br>H3K9AC |
| chr22_32474782_C_T_b38* | Yes | Yes | CTCF,<br>H3K27AC,<br>NEUROD1,<br>NR2F2,<br>RAD21,<br>ASH2L,<br>CC2D1A,<br>H3K9AC,<br>NONO,<br>TRIM28 | CTCF,<br>H3K4ME3,<br>POL2,<br>RAD21,<br>SOX8, SP1,<br>ATF7,<br>H3K9AC,<br>ZBTB7B |
| chr22_32474819_C_T_b38** | No | Yes | NA |  |

## Discussion

Here, we present an analysis of allele-biased binding across 93 transcription factors, identifying thousands of variants that show biased binding. We identified a threshold for reproducibility that provides confidence to our calls both within a single donor and across multiple donors, controlling for a wide variety of biological and technical variables. By linking to allele-biased expression of nearby genes, we also relate variation that impacts TF binding directly to effects on gene regulation.

We found that TF-biased variants are prevalent in distal and proximal enhancer regions as well as in promoter regions. This highlights that these variants occur in regions known to play major roles in gene expression. This, combined with the many cases of allele-biased binding within eQTLs, shows a potential mechanism of eQTLs, and may lead to insights of disease mechanisms (Musunuru et al. 2010). Because a majority (>80%) of the allele-biased variants fall outside of called peaks for the biased TFs, this also stresses the importance of TF binding outside of peaks that have measurable impact, as noted in previous studies (Lun and Smyth 2016; Hiatt et al. 2023).

We found that rare variation (MAF < 0.1%) is enriched, relative to common variation, for TF-binding impacts (**Figure 3B**), suggesting that there was purifying selection against such variation in general. For common variants that do impact binding, neither the ancestral nor derived alleles tend to be favored (**Supplemental Figure 7**). In contrast, among rare variants (MAF < 0.1%), there is a bias in favor of common alleles over rare alleles, whether the common allele is derived or ancestral. This suggests that new mutations more often disrupt, rather than enhance, TF binding. Still, the fact that TF-variant bias can sometimes prefer the novel allele even in rare variants is consistent with models of *de novo* motif formation and gene birth (Behrens and Vingron 2010; Schlötterer 2015; Carvunis et al. 2012; Iyengar and Bornberg-Bauer 2023; Ruiz-Orera et al. 2015; Papadopoulos et al. 2021), which have suggested that few changes need to be made to a given sequence to form a novel TF motif, and that this formation plays a crucial role in sampling of transcriptional regulatory space. This is emphasized by the fact that we observe a small subset of derived alleles that have become common in the population (DAF>=0.999), and are favored by TFs (**Supplemental Figure 7**).

As has been previously observed (Abramov et al. 2021), in cases of TF-biased variants, there is a general preference for a given TF to favor the allele with a stronger presence of its motif, and TF read-depth measurements confirm a correlation between the degree of motif disruption and total read-depth for variants within motifs (**Figure 3D**). Beyond demonstrating the general nature of this phenomenon, it can be combined with known eQTLs in our dataset to facilitate fine-mapping and mechanistic hypothesis generation. For example, we highlight a case of a variant in an eQTL that displays allele biased binding in our dataset and specifically disrupts a motif for that TF (**Figure 5**), affecting regulation of a gene that is relevant to neurodegenerative and neuropsychiatric traits. The results from this analysis yielded TF-biased variation linked to 9,748 GTEx eQTLs, providing a rich resource for future fine-mapping efforts.

We also found that many sites of allele-biased binding represent coordinated multi-factor effects. For example, 48.1% of sites that associate with altered binding of one TF influence binding of one or more additional TFs (**Supplemental Tables 1-2**). Similarly, approximately half of variants with TF-binding bias also have altered histone marks and POL2 binding, consistent with the expected relationships between TF binding and general recruitment of transcriptional machinery. Finally, we found that approximately 30% of TF-allele-biased variants in our data impacted cohesion complex members (**Supplemental Figure 9**). This suggests that genetic variants which alter three-dimensional genome interactions are a major contributor to gene expression variation in the population.

Overall, our study provides a resource of allele-biased variants that are experimentally validated to impact TF binding in a biologically relevant context. These results will further our understanding of how alteration in DNA sequence translates to changes in biological function, particularly in relation to analyses of gene regulation in the human brain.

## Methods

### Whole Genome Sequencing and variant calling

We extracted high molecular weight DNA from approximately 20 mg cortex tissue from each donor using the MagAttract HMW DNA kit (Qiagen 67563). We prepared linked read libraries using the Chromium Genome Reagent Kit v2 following the protocol provided by 10x Genomics. We processed sequence reads using the longranger software suite from 10x Genomics. We identified variants by aligning to a 10x Genomics-provided, longranger-enabled hg38 reference (version 2.1.0) using longranger wgs v2.2.2. We called variants using GATK 3.8-1-0-gf15c1c3ef via the –vcmode gatk option in the longranger wgs workflow (Loupe et al. 2023).

### Genome Construction

We constructed graph genomes using the vg toolkit version 1.20, available at https://github.com/vgteam/vg (Garrison et al. 2018). The “construct” command was used with the hg38 genome and all phased variants which passed quality metrics. We then pruned the graph using the “prune” command with default parameters. We produced the gbwt index using the “index” command with default parameters, and the gcsa index was created using the parameters: -X 3 -Z 4000 -p -k 11.

We also constructed linear FASTA sequences for comparisons of linear and graph genome reference allele bias. We identified variants which were within 1 full read length (100 bp) of one another, and on the same phase. We identified regions based on such nearby, in-phase variants, and constructed a fasta file containing, for each such region, one entry for either haplotype. In these haplotypes, 78.6% of regions contain only a single variant, while 96.2% had no more than 2 variants.

### RNA-seq

We performed RNA-seq for each of the nine brain regions as outlined in Loupe *et al*. 2023.

### ChIP-seq experiments

We peformed ChIP-seq experiments with 93 TFs and five histone marks in nine distinct brain regions, for a total of 1,028 experiments. Full methods for production of ChIP-seq reads are presented in (Loupe et al. 2023).

### Peak Calling

We called peaks according to the ENCODE Consortium’s standard pipeline, using experiments from donors as replicates, as described in (Loupe et al. 2023).

### Read Mapping and processing

For traditional read mapping, we used bowtie2 (Langmead and Salzberg 2012) with default settings to map to the human hg38 genome.

For graph genome mapping, we used the vg map command with arguments -A -K -M 3. The vg surject command was used to create sam and bam file formats for determining allele bias. The samtools package (Danecek et al. 2021) was used for sorting and filtering by quality. Picard was used for filtering duplicates.

Once reads were mapped and filtered, we identified and separated out only those reads that overlapped with a heterozygous variant using custom R code. In brief, for a read mapped to a heterozygous region, we determined the minimum string distance, i.e. greatest sequence similarity, between the read and each of the two haplotypes, and assigned the read to the haplotype that was most similar to the read’s sequence. In cases where the minimum string distance between the two haplotypes was equal, we assigned half a read to each sequence, leading to a more conservative binomial test for allelic bias.

### Identification of Allele Bias

For a given ChIP-seq or RNA-seq dataset, after mapping, we identified those heterozygous regions with at least six total reads (the minimum number of reads for a binomial test to be nominally significant at p<=0.05 if all reads map to a single haplotype). After assigning a number of reads to each haplotype, we performed a two-sided binomial test for each haplotype for each ChIP-seq dataset.

After assessing the consistency of allelic biases across brain regions and donors, we summed the number of reads assigned to each haplotype for a given TF across all tested brain regions within a single donor, and a two-sided binomial test was performed for each haplotype for each TF with at least six reads when combined across tissue samples, the minimum number of reads required for a significant binomial test p-value at a 0.05 cutoff. For some analyses, we restricted to cases of at least 11 reads total, the minimum number of reads required for a significant binomial test p-value at a 0.001 cutoff. In all analyses, we removed variants that showed apparent allele bias in control input DNA for the summed dataset in the respective donor. These variants are included in **Supplemental Tables 1 and 2**.

### VEP Annotations and Derived Allele Frequency

We annotated vcf files using the following command:

vep -i 5397-JL-0002_phased_variants.vcf.gz --config vep108.ini --vcf -o 5397-JL-0002_annotated.vcf.gz

The config file is provided in the Supplemental_Code.zip file as vep108.ini. VEP engine and cache version 108 (McLaren et al. 2016) was used with a GRCH38 fasta file. We used a merged transcript set of Ensemble (Cunningham et al. 2022) and RefSeq (O’Leary et al. 2016). Custom annotations were Gnomad (Chen et al. 2022) allele frequency using v 3.1.1, Bravo topmed allele frequency freeze 8 (Taliun et al. 2021), GRCh38 GERP scores (as distributed with CADD v1.6), and CADD v1.6 scores (Rentzsch et al. 2019).

We treated each variant in the haplotype separately in the rare cases where a single haplotype region contained multiple variants with different Derived Allele Frequencies and that haplotype region showed TF binding bias,.

### GTEx Data and identification of RNA allele bias

We downloaded GTEx variants on June 30^th^, 2023 from https://storage.googleapis.com/gtex_analysis_v8/single_tissue_qtl_data/GTEx_Analysis_v8_eQTL.tar

For a given eQTL variant in our data, we determined whether or not there was a phased heterozygous variant within the appropriate gene body in our data, as this was necessary for physically linking the TF-allelic bias to RNA allelic bias. In such cases, we determined the variant in the gene body which was on the same allele as each of the two haplotypes of the heterozygous variant in the GTEx dataset. We calculated significant bias as discussed above, and we calculated effect size as:

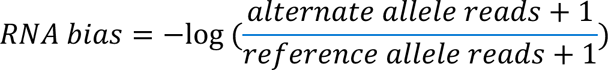

### GREAT Gene Ontology Analysis

We performed GREAT analysis using the rGREAT (v2.1.8) package (https://www.bioconductor.org/packages/release/bioc/html/rGREAT.html) (Gu and Hübschmann 2023). We associated genomic ranges with genes using the basal plus extension method (5kb upstream, 1kb downstream, 500kb max extension). We calculated enrichment for GO Biological Process terms within GREAT with background regions set as the union of all ChIP-seq peaks with heterozygous variation that did not show evidence of allele-biased binding.

### Data Analysis

We performed data analysis using R version and 4.1.0 (2010), as noted in appropriate scripts.

### cCRE catalog

We downloaded the V4 cCRE human dataset from the ENCODE Portal under accession ENCSR800VNX.

## Data Access

All code used for these analyses is available via GitHub at https://github.com/bmoyers/BrainTF_Allele_Biased_Binding, and is also supplied as Supplemental_Code.zip. These data and the accompanying analyses will serve as a resource to understand genome regulation in psychiatric diseases and are publicly available through the PsychENCODE Consortium and available for download at the following link: https://doi.org/10.7303/syn4921369.

## Acknowledgements

We thank the brain donors and their families without whom this research would not have been possible. We thank members of the Myers, Cochran, G. Cooper and S. Cooper labs for many fruitful discussions and input.

